# Long-range quantum coherence of the photosystem 2 complexes in living cyanobacteria

**DOI:** 10.1101/2019.12.13.875344

**Authors:** T. Rammler, F. Wackenhut, S. zur Oven-Krockhaus, J. Rapp, K. Forchhammer, K. Harter, A. J. Meixner

## Abstract

The first step in photosynthesis is an extremely efficient energy transfer mechanism, which is difficult to be explained by classical short-range energy migration (“hopping”) and led to the debate to which extent quantum coherence is involved in the energy transfer between the photosynthetic pigments. Embedding living cyanobacteria between the mirrors of an optical microresonator and using low intensity white light irradiation we observe vacuum Rabi splitting in the transmission and fluorescence spectra as a result of strong light matter coupling of the chlorophyll and the resonator modes. The Rabi-splitting scales with the number of chlorophyll a pigments involved in coherent coupling indicating forming a polaritonic state which is delocalized over the entire cyanobacterial thylakoid system, down to the single photon level. Our data provide evidence that a delocalized polaritonic state is the basis of the extremely high energy transfer efficiency under natural conditions.

## Introduction

In photosynthesis, light energy is absorbed and converted into relatively stable chemical products by membrane-integral pigment-protein complexes called photosystems for long-term chemical energy storage.^1^ Photosynthetic complexes are optimized to capture photons from solar light and transmit the excitation energy from peripheral pigments to the photosynthetic reaction center with an extremely high efficiency (close to 100 % ^2^). They consist of a collection of pigment molecules, such as chlorophylls and carotenoids, that are arranged by a protein scaffold in a way that near-field dipole coupling is possible.^2^ When interacting with light they no longer act as independent excited molecules, but coupling between them results in collective excitations called excitons, whose wave function extends over several chromophore units.^3 4^ Electronic interaction with the local environment tunes individual pigment excitation levels to form an energetic ladder from higher energy at the periphery to lower energy near the reaction center, often described semi-classically as “sequential hopping” ^5^ of excitation energy. On their way to the reaction center, the excitons loose part of their energy to the vibrational modes of the protein complex that serve as a thermal bath. The transfer efficiency depends on the distance, spectral overlap of the emission and absorption spectrum and relative orientation of the pigment molecules to each other. However, structural analyses of photosynthetic complexes ^6^ show that none of these factors is optimal for the observed highly efficient energy transfer. Finally, the transfer competes continuously with fluorescence emission and non-radiative deactivation of the excited chlorophylls.

The observation of oscillatory intensity modulations of ultrafast photon echoes from isolated photosynthetic complexes of *Chlorobium* at cryogenic temperatures and under almost physiological conditions has drawn enormous attention and led to the hypothesis that quantum coherence could be a possible explanation for this efficient energy transfer. ^7 3 8 9 10 11^ Recent investigations have revealed that both electronic and vibrational coherences are involved in primary energy transfer in bacterial reaction centers ^12 13^. However, the physiological relevance of photosynthesis-related quantum coherence has rarely been studied in living photosynthetic organisms. Furthermore, excitation in ultrafast time-resolved lasers spectroscopy is pulsed and coherent, while irradiation in nature occurs continuously over the course of minutes to hours by incoherent photons. As a consequence, the energy transfer in the photosynthetic machinery must operate on the basis of independent single photons. Additionally, it is not clear whether the occurrence of quantum coherence might even be a prerequisite for the function of the photosynthetic machinery and provides a selection advantage in the development of photosynthetic organisms.

To address this open question, we have enclosed living cyanobacteria (*Synechococcus elongates*) in the confined electromagnetic field of an optical microresonator to probe their optical properties *in vivo*. Using low intensity white light irradiation, we show that long-range coherent excitonic coupling between the chlorophyll a pigments of PS2 via the formation of a polaritonic state, delocalized over an entire cyanobacterium, is probably one reason for the very efficient photosynthetic energy transfer not only in cyanobacteria but also related plant chloroplasts.

## Results

To study possible long-range quantum coherence effects in the photosynthesis of living organisms at ambient conditions, we embedded cells of *S. elongatus* (strain PCC 7942) in an optical microcavity. In contrast to sulfur bacteria ^14 3^, *S. elongatus* performs oxygenic photosynthesis that is more related to the photosynthesis of plant chloroplasts and, based on the available data ^15^, is a particularly well-suited photosynthetic model for the study of quantum physical processes *in vivo* at physiological conditions.

Our Fabry-Pérot optical microresonator (quality factor, Q = 98) consists of two partially transparent mirrors (Fig. 1A). Their distance can be precisely adjusted with a piezo actuator to control the resonance condition of the microcavity. Compared to previous works ^14^, we have chosen silver mirrors with a large layer thickness to achieve a stronger interaction between the microcavity and the cyanobacteria. More details about the experimental set up are given in the supporting information. Transmission spectra were acquired from below via a high numerical aperture (NA) objective lens (NA = 1.4), while the microcavity was irradiated by a continuously emitting white light source from above (Fig. 1A). Additionally, we irradiated the sample with a laser from below to detect strong coupling in the emission spectrum of individual bacteria, which is made possible by the small focal spot size of the high NA objective. As shown in Fig. 1B, the *in vivo* absorption (blue line) shows the typical chlorophyll a maximum at around 680 *nm* due to a red-shift of protein-bound chlorophyll. ^17^ The emission spectrum at 440 *nm* excitation reveals again a maximum at around 680 *nm* which is assigned to the PS2 complexes and phycobilisome terminal emitters (APC680). ^18 15^ The cavity resonance can be tuned across the absorption and emission maximum (see Fig. S1), allowing efficient optical coupling between the cavity modes and the cyanobacteria. Remarkably, the absorption and emission spectra show a significant overlap (Fig. 1B) demonstrating that the cyanobacteria are able to reabsorb their own emitted light. This photophysical feature of *S. elongatus* drastically increases the chance that its photosynthetic pigments can strongly couple to an optical field confined in a microcavity.

**Fig. 1:**
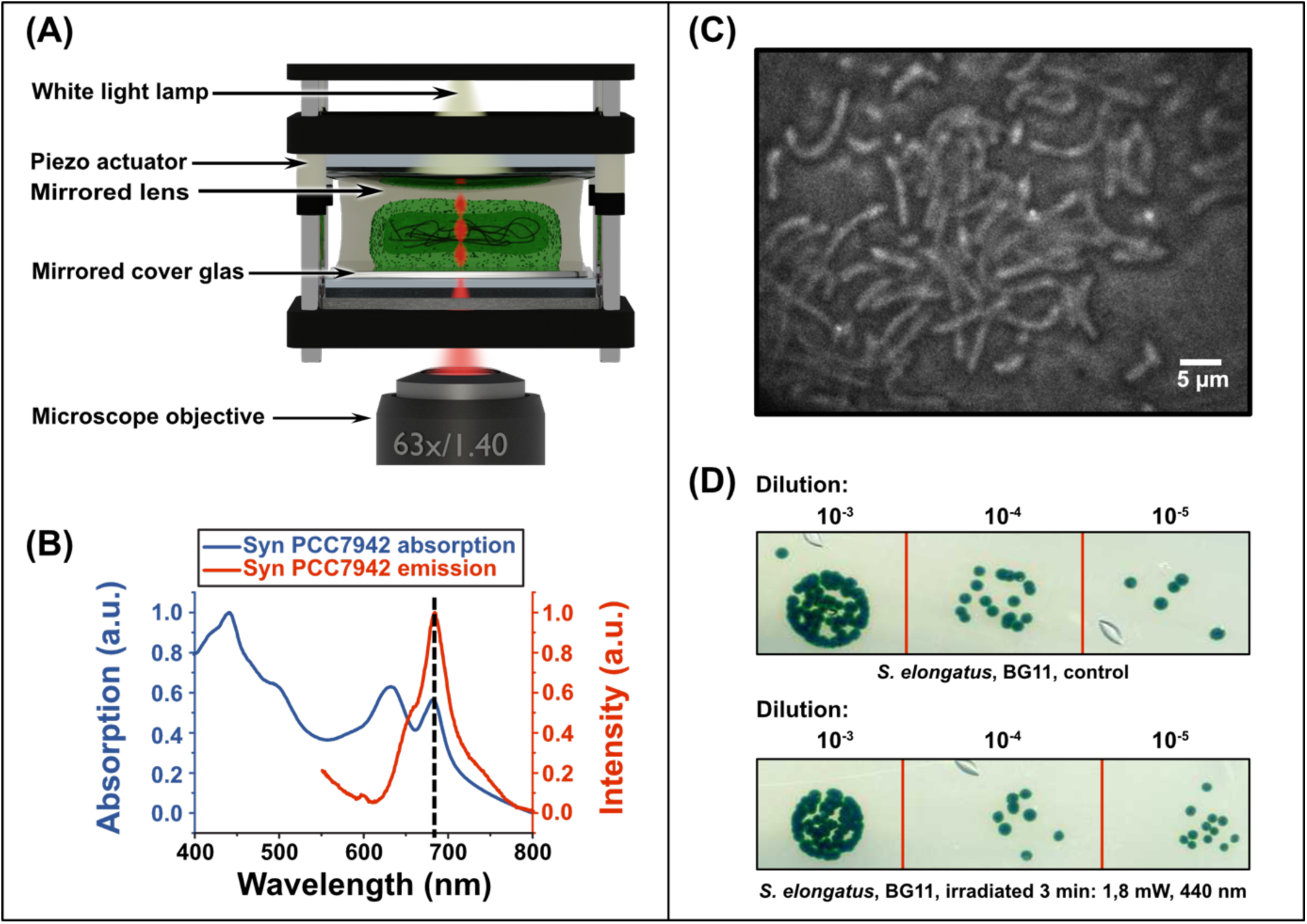
The spectral properties of the photosynthetic pigments of *S. elongatus* cells and a Fabry-Pérot microresonator. The survival of *S. elongatus* cells is not impaired by the light conditions prevailing in the microcavity. (**A**) Scheme of the Fabry-Pérot microcavity, which consists of two partially transparent mirrors. The distance between the mirrors can be fine-tuned with piezoelectric actuators. Due to constructive and destructive interference, only wavelengths fulfilling the resonance condition of the cavity are transmitted. The bacteria are placed in an agarose matrix inside the cavity. (**B**) Normalized absorption (blue) and fluorescence (red) spectra (*λ*_*ex*_ = 440 *nm*) of *S. elongatus* cells located inside the microcavity. The dashed black line indicates the wavelength where the bacteria emit and absorb photons of the same wavelength. (**C**) Light microscopy image of *S. elongatus* cells inside the microcavity. (**D**) Spot assay ^16^ of *S. elongatus* cells in BG11 medium. Top: Non-irradiated control in a dilution series (1:10), initial concentration: *OD*_750_ = 0.5. Below: Irradiated sample in a dilution series (1:10), initial concentration: *OD*_750_ = 0.5. The bacteria were irradiated with a laser before preparation of a spot assay. The irradiation conditions were comparable to those in the microcavity. The comparable growth rate indicates a negligible impact of the typical irradiation during the experiments.

The survival rate was assayed to examine the possible impact of the laser irradiation on the cyanobacteria embedded in the microcavity (Fig. 1C). Since only a single bacterium is exposed to the focused laser irradiation inside the cavity at a time, which cannot be isolated after the experiment, we have designed an assay to analyze comparable irradiation conditions by embedding a cyanobacterial culture in a low-melting agarose matrix outside the cavity. The cyanobacteria were then exposed to light conditions (440 *nm*, 1.8 *mW*, 3 *min*) similar to those prevailing in the cavity, while a non-irradiated culture served as a control and the survivability was analyzed by a spot assay (see supporting information for details). No growth difference between the irradiated and non-irradiated sample was observed (Fig 1D), indicating that the light conditions in the microcavity have no discernible impact on the cyanobacterial survivability.

To determine, whether the influence of the microcavity on the cyanobacterial photosynthetic system is detectable *in vivo*, fluorescence lifetimes (FLT) ^19^ of the photosynthetic pigments in single bacteria were acquired.

The light-harvesting pigments of the photosynthetic complex serve to rapidly and efficiently transfer light energy from the peripheral pigments to the reaction center, therefore, the fluorescence signal of cyanobacteria is weak. Transfer and trapping of the excitation energy in the photosystem leads to a fast non-exponential fluorescence decay, which can be observed from live cyanobacteria with a wide distribution of fluorescence life-time (FLT) components from short ones in the low and mid picosecond range and slow components in the low nanosecond range.^20^ The spontaneous emission rate of a chromophore can be increased or decreased by placing it in a microcavity and tuning it in-resonance or off-resonance with the chromophore emission. This is known as Purcell effect ^21 22^, leading to shorter or longer FLTs, respectively. The long lived FLT component originates from particularly those photosynthetic pigments, where the excitation energy is trapped in an emitting state and can therefore be analyzed *in vivo* for three cases: (i) free space (outside of the cavity), (ii) inside the cavity in off-resonance mode and (iii) inside the cavity in resonance mode. Short laser pulses (*λ*_*ex*_ = 440 *nm*) with pulse durations of less than 80 *ps* and a pulse rate of 80 *MHz* were used. The pigments irradiated in free space (i) reveal a FLT value of *τ*_*I*_ = (0.26 ± 0.006) *ns* ((i) in Fig. 2) and a slightly larger value of *τ*_*I*_ = (0.29 ± 0.016) *ns* is recorded in the off-resonant microcavity ((ii) in Fig. 2). In contrast, the FLT in the resonant microcavity (iii) decreased to *τ*_*I*_ = (0.16 ± 0.006) *ns* and was significantly shorter compared with the data obtained in free space or in the off-resonant cavity. This result is consistent with the Purcell effect ^21^ of isolated chromophores and demonstrates that the microcavity has a noticeable impact on the photosynthetic processes in single living cyanobacteria.

**Fig. 2:**
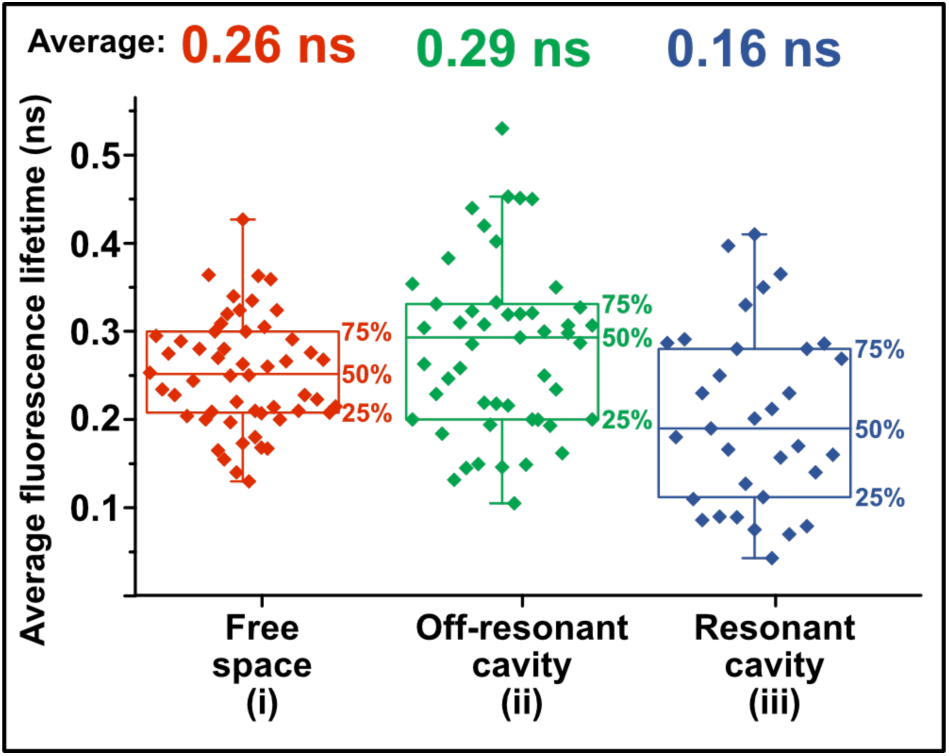
The fluorescence lifetime (FLT) of *S. elongatus* photosynthetic pigments is influenced *in vivo* by the microcavity. The bacteria were embedded in low-melting agarose and irradiated with short laser pulses (*λ*_*ex*_ = 440 *nm*) with a duration of less than 80 *ps* and a repetition rate of 80 *MHz*. The average intensity-weighted FLTs were recorded in free space ((i), red), inside the off-resonance cavity ((ii), green) or inside the cavity in resonance with the light emission of the cyanobacteria ((iii), blue). A two-tailed t-test confirms a significant difference between the fluorescence lifetimes for the off-resonant (and without) cavity and the resonant cavity, *p* = 2.05 · 10^−5^, (*p* = 3.20 · 10^−6^).

In general, the interaction of a quantum system with the optical field in a microcavity can be separated in the weak and strong coupling regime. In the weak coupling regime, the individual damping constants of the cavity and the photosynthetic pigments are larger than the coupling constant. In this case, the microcavity only influences the spontaneous emission rate *via* the Purcell effect as observed in the FLT analysis (Fig. 2). However, if the coupling constant exceeds the individual damping constants, the energy of a photon can be coherently cycled back and forth between the oscillating electromagnetic field in the microcavity and the induced polarization formed by a large number of coherent electronically excited chromophores enclosed between the cavity mirrors before it escapes from the cavity; this condition reflects the strong coupling regime ^23^. The energy of the photon, which is dispersed in the whole mode volume and shared between the cavity mode and the polarization, is described in quantum electrodynamics as a hybrid light-matter state or polariton. ^24 25 26^ In our case, these so-called polaritonic modes are a coherent superposition of the cavity mode and the electronically excited state of the photosynthetic pigments (chlorophyll a in PS2). As shown in Fig. S2, the back and forth cycling of the photon energy between the electromagnetic field in the microcavity and the polarization in the time domain leads to a splitting in the spectral domain that manifests itself as a double-peaked cavity transmission spectrum with a peak separation referred to as vacuum Rabi splitting, as schematically illustrated in Fig. 3A/B.

**Fig. 3:**
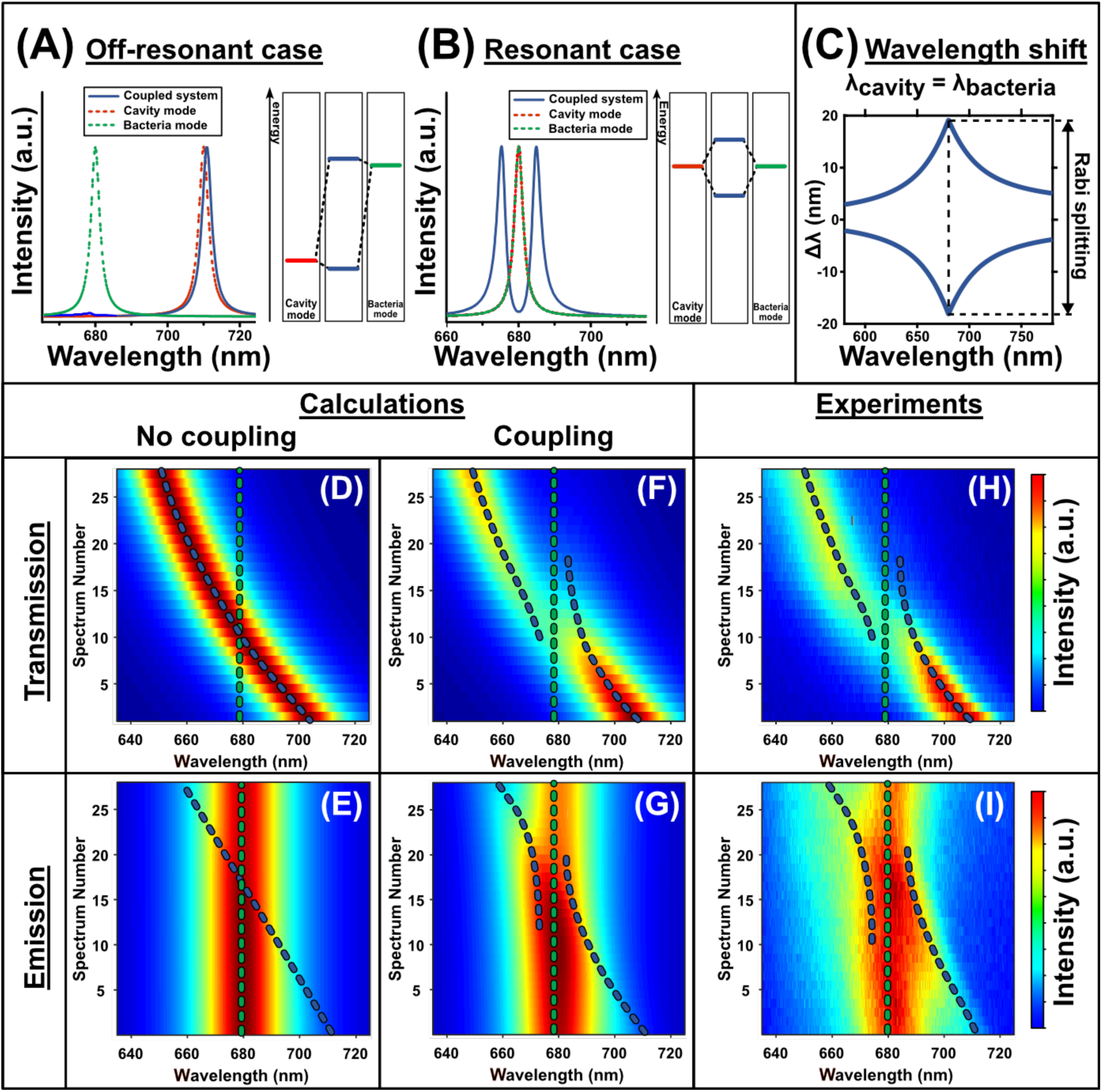
Strong coupling between a microcavity and the photosynthetic machinery of living cyanobacteria. (**A**) The dashed green and red spectra represent the uncoupled bacteria emission/cavity mode. The blue spectrum illustrates the cavity transmission spectrum for the non-resonant but coupled case and is similar to the uncoupled system. The graph on the right illustrates the corresponding energy level scheme. (**B**) Illustration of the resonant case, where the cavity mode is spectrally overlapping with the bacteria emission, and two polaritonic modes (blue lines) are clearly visible in the double-peaked cavity transmission spectrum. (**C**) Spectral shift *Δλ* of the coupled modes relative to the uncoupled ones. The largest splitting, i.e. vacuum Rabi splitting, is observed when the cavity and bacteria are in resonance. (**D**)/(**E**) Simulated cavity transmission/ bacteria emission spectra without coupling as a function of the decreasing mirror distance (indicated by the spectrum number). The dashed green and blue lines are the spectral position of the uncoupled bacteria emission/cavity resonance, respectively. No anti-crossing can be observed when the cavity mode is tuned across the bacteria emission. (**F**)/(**G**) Simulated cavity transmission/bacteria emission spectra including strong coupling between the cavity mode and the bacteria emission. Strong coupling is visible in (F)/(G) by the anti-crossing dispersion, when the cavity mode is close to the emission of the bacteria. (**H**)/(**I**) Experimental cavity transmission/bacteria emission spectrum. Strong coupling can be observed in **(H)/(I)** by the anti-crossing dispersion and is in perfect agreement with the simulation in (F)/(G).

To study vacuum Rabi splitting, due to a polaritonic mode in the photosynthetic machinery of a living cyanobacterium and the optical field in the microcavity, we simulated and experimentally investigated the dispersive behavior of the coupled system.

The dashed lines in Fig. 3A/B illustrate the simulated uncoupled photosystem emission of the cyanobacteria (green) and the cavity mode (red). Fig. 3A illustrates the case when there is no spectral overlap between them. The transmission spectrum of such a coupled, but off-resonant system is similar to that of the uncoupled cavity mode. Conversely, when the cavity mode approaches the spectral position of the chlorophyll a emission, a splitting into two polaritonic modes is visible (Fig. 3B, blue line). Fig. 3C represents the simulated spectral shift *Δλ* of the coupled modes relative to the uncoupled modes. The shift caused by strong coupling is largest when the cavity is in resonance with the chlorophyll a emission, leading to a symmetric double-peaked cavity transmission spectrum. The occurrence of such a spectral gap between the two polaritonic modes is called vacuum Rabi splitting. Mathematically, such a coupled system can be modeled by two coupled damped harmonic oscillators, as described in the supporting information or in ^27^. First, we want to illustrate in Fig. 3D/E the results of the calculation without coupling (*κ* = 0 *eV*) between the cavity mode and the chlorophyll a emission. Each line in Fig. 3D represents a cavity transmission spectrum, as indicated by the spectrum number, and its intensity given by the color map. In this simulation, the cavity length gradually increased from top to bottom, leading to a spectral red shift of the cavity resonance. The dashed lines in Fig. 3D/E represent the spectral position of the uncoupled chlorophyll a emission and the cavity mode, respectively. In the absence of strong coupling, no splitting is observed, even when both modes were tuned to the same resonance wavelength; the chlorophyll a emission was only influenced by the Purcell effect (Fig. 3E). This changes with strong coupling between the cavity mode and the chlorophyll a pigments, with a calculated coupling constant of *κ* = 0.14 *eV* in Fig. 3F/G. The calculated cavity transmission spectra in Fig. 3F show a clear anti-crossing behavior when the cavity resonance approaches the spectral position of the chlorophyll a emission at 680 *nm*. In the calculated emission spectra in Fig. 3G, the mode splitting is less obvious because it is obscured by the spectrally broad fluorescence background of chlorophyll a pigments that do not contribute to the polarization. The uncoupled pigments have their electronic transition dipole moments oriented perpendicular to the polarization and constitute about 2/3 of the total number of pigments. By comparing the simulations in Fig. 3D/E with Fig. 3F/G, it is possible to experimentally distinguish between no/weak and strong coupling in the microcavity-cyanobacteria system. Notably, the experimental white light transmission spectra derived from the living cyanobacteria show a clear anti-crossing behavior when the cavity resonance is tuned over the chlorophyll a emission at 680 *nm*. (Fig. 3H). In the emission spectra (Fig. 3I) the splitting is less obvious since it is composed of two types of photons, those that participate in the strong coupling process with the cavity mode and those that escape from the resonator without coupling due to the Purcell effect. The experimental results fit perfectly to the calculated spectra in Fig. 3F/G and prove that strong coupling between the microcavity and the chlorophyll a pigments is achievable in living cyanobacteria.

According to the Jaynes-Cummings model the energy splitting *ΔE* is given by equation (1) and is proportional to the square root of the number *n* of fluorophores that coherently couple to the cavity mode with a coupling constant *g*_0_. ^23^

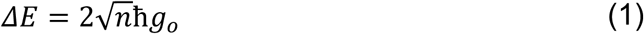

Therefore, the splitting energy should decrease when the number of fluorophores is reduced. This can indeed be achieved in a living bacterium by photobleaching a fraction of the functional chlorophyll a pigments by increasing the laser intensity by a factor of 100 as compared with the previous experiments.

As shown in Fig. 4A at the beginning the cavity mode at *λ* = 680 *nm* has a spectral dip at the center due to vacuum Rabi splitting and as the photobleaching of the pigment molecules proceeds (Fig. 4A, blue dashed line) the energy splitting between the two peaks reduces and disappears. In contrast, at the same time for the cavity mode at *λ* = 546 *nm* which has no coupling to the chlorophyll a pigments no changes in intensity or spectral position are visible. This is further illustrated in Fig. 4B, where the first (blue line) and the last (red line) spectrum of the spectral series in Fig. 4A are shown.

**Fig. 4:**
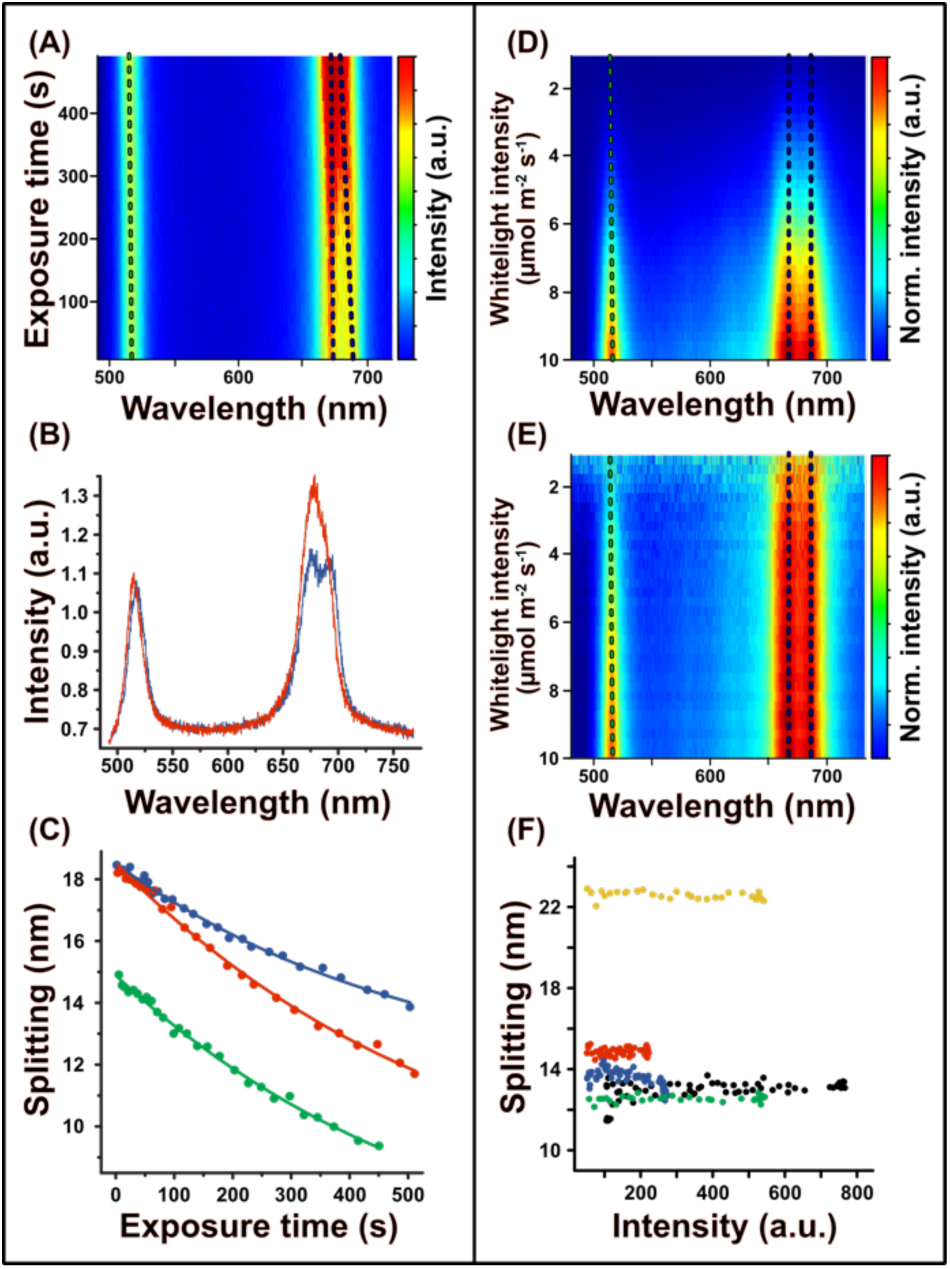
Reducing the number of pigments in a cyanobacterium by photobleaching reduces vacuum Rabi splitting and shows that pigments all over a bacterium are coherently coupled. (**A**) Cavity transmission spectra with two cavity modes as a function of the exposure time. One mode shows vacuum Rabi splitting (dashed blue line), while the other is not coupled (dashed green line). The splitting energy, and thus the coupling, is reduced by the continuous irradiation and photobleaching the pigments of the bacterium. (**B**) First (blue line, *t* = 0 *s*) and last (red line, *t* = 500 *s*) spectrum of A. (**C**) Rabi splitting between the two intensity maxima around 680 *nm* as a function of the exposure time. Three different, individual bacteria (red, green and blue) in the cavity show the decrease of the vacuum Rabi splitting with increasing bleaching of the pigments. (**D**) Cavity transmission spectra with two resonances as a function of the illumination intensity of the white light lamp of 10 µ*mol* photons s ^−1^ m ^−2^ to 1 µ*mol* photons s ^−1^ m ^−2^ (corresponding to 28 – 2.8 *mWcm* ^−2^ at 680 *nm*) from bottom to top. The coupling remains constant while the intensity of the white light lamp is reduced. (**E**) Intensity normalized version of (D), where each spectrum is normalized to its maximum intensity, to better visualize the constant splitting. (**F**) Splitting as a function of the white light intensity. At low illumination intensity, five different, individual bacteria in the cavity show that the vacuum Rabi splitting is independent of the light intensity.

The number of molecules (*n* in Eq. 1) decreases exponentially by photobleaching as tested at three different locations in the cavity, resulting in a decreased splitting of the coupled modes, which can be fitted to the square root of an exponential decay (Fig. 4C). These results demonstrate that the extent of the Rabi splitting depends on the number of pigments effectively participating in polaritonic coupling to the optical mode throughout the entire focal volume.

In general, the photosynthetic efficiency would benefit enormously when the coherence between the light harvesting pigments and the cavity is independent of the light intensity. To reveal the light intensity dependence, white-light transmission spectra were acquired with different excitation intensities of 10 µ*mol* photons s^−1^ m^−2^ down to 1 µ*mol* photons s^−1^ m^−2^ (corresponding to 28.0 – 2.8 *mWcm*^−2^ measured at 680 *nm*) in single living cyanobacteria as shown in Fig 4D/E, where the y-axis corresponds to different excitation intensities. The resonance mode at *λ* = 680 *nm*, which is strongly coupled to chlorophyll a, shows a Rabi splitting which remains constant with decreasing white light irradiation intensity (Fig. 4D). This is even more obvious in the normalized spectra (Fig. 4E), where each spectrum along the y axis is normalized to its maximum intensity. This constant Rabi splitting is observed for different individual cyanobacteria in the sample, as shown in Fig. 4F by plotting the Rabi splitting against the intensity of the cavity mode. As a consequence, since the photons used for white-light illumination are completely incoherent, strong coupling must occur even at very low light intensity; or in other words, one resonant photon is already sufficient to induce a polaritonic state between the microcavity and the chlorophyll a pigments *in vivo*.

## Discussion

The emission and transmission spectra presented here demonstrate undoubtedly the existence of an *in vivo* coherent energy exchange between the microcavity and the cyanobacterial photosynthetic light harvesting machinery. Photo-bleaching experiments confirm that the microcavity can couple to about 4.8 · 10^5^ chlorophyll a molecules (calculated from the measured splitting and assuming a chlorophyll a transition dipole moment: 5.39 *D* @ *n* = 1.34 ^28^, see supporting information for calculation) at the same time to form a polaritonic state delocalized over the entire thylakoid system of a single cyanobacterium (Fig. 5A). For natural illumination conditions, the semi-classical model of energy “hopping” (Fig. 5B) in the photosynthetic light-harvesting machinery should therefore be expanded to a delocalized wave-like energy transfer (red glow in Fig. 5C). Additionally, the vacuum Rabi splitting is independent of the number of photons, indicating that it works at very low light intensities and for the formation of a polaritonic state one single photon is sufficient. The cell-encompassing delocalization of the polaritonic state guarantees that the photosynthetic reaction centers are optimally supplied with photons and allow growth and survival of the cyanobacteria even in a low-light environment.

**Fig. 5:**
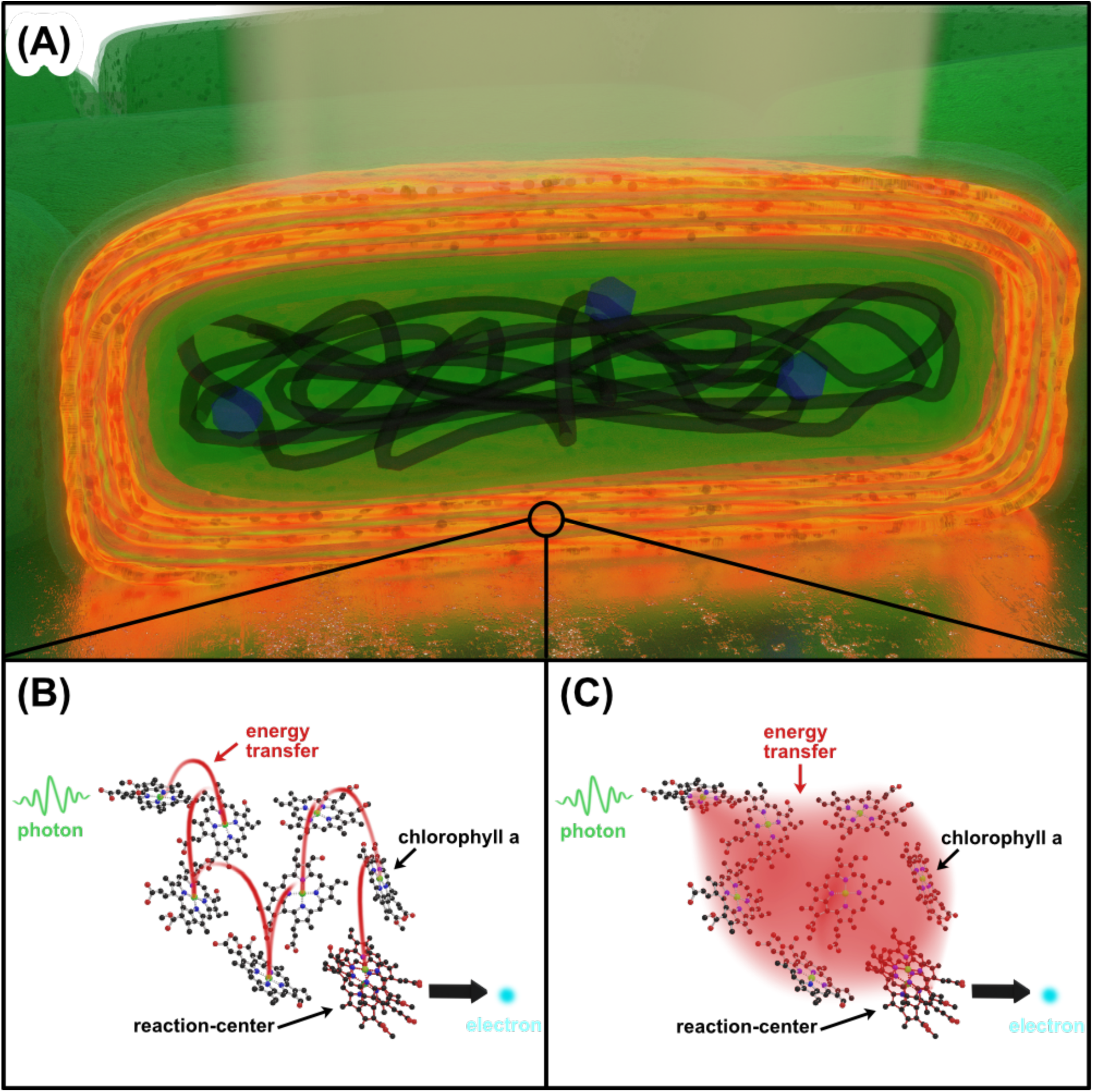
Models of energy transfer during photosynthesis. **(A)** Illustration of the energy delocalization over the thylakoid system. (**B**) Classical description: after absorption of a photon (green wavy arrow), the energy transfer from the antenna pigments (chlorophyll a molecules) to the reaction center (chlorophyll a dimer) is described by incoherent energy “hopping” (red energy transfer). (**C**) A collective coherent state between a photon and the antenna pigments generates a coherent energy transfer, being the cause for the very high energy transfer efficiency (close to 100 %).^30^

The establishment of a long-range polaritonic state requires a complex and dynamic spatial and structural organization of the photosynthetic complexes that compensate or make use of the thermodynamic fluctuations occurring in the natural environment. It can therefore be assumed that the observed cell-encompassing quantum coherence is not merely a byproduct of the evolution of photosynthesis. On the contrary, evolution has rather selected for this process of coherence to optimize photosynthesis to its highest efficiency and makes it very likely that it also occurs in plant chloroplasts, since cyanobacteria are their precursors. As a consequence of our validation of quantum coherence in the photosynthetic machinery of living cyanobacteria at physiological conditions, we believe that the investigation of various other biological processes, which are difficult to be explained by classical thermodynamics, may reveal significant dependencies on quantum electrodynamics effects.^29 2^

## Methods

### Preparation of cavity mirrors

The mirrors were produced by evaporating a 3 *nm* thick chromium layer on a glass surface serving as an adhesion layer for the following silver layer, which has a thickness of 30 *nm* or 60 *nm* for the lower and upper mirror, respectively. Since silver is bactericidal and very susceptible to damage and oxidation, it was coated with a gold layer (5 *nm*) and an *SiO*_2_ layer (20 *nm*).^31^ These layer thicknesses result in a microcavity with a quality factor of *Q* = 98. The microcavity was assembled in a custom-built holder with piezo actuators and mounted on a stage scanning confocal microscope for the collection of both white light transmission and fluorescence spectra from the same spatial position.

### Light intensity measurements

The light intensity was measured with a Li-Cor Li-189 radiometer from Heinz Walz GmbH (Germany).

### Bacterial cultivation conditions

*Synechococcus elongatus* PCC 7492 cells were cultivated under photoautotrophic conditions with continuous illumination at around 30 µ*mol* photons *s* ^−1^ *m* ^−2^ (Lumilux de Lux, Daylight, Osram) at 28 °*C*. The cultures were grown in 100 *mL* Erlenmeyer flasks, filled with 40 *mL* BG11 ^32^ medium, supplemented with 5 *mM NaHCO*_3_ and shaken at 120 − 130 *rpm*.

### The survivability after laser irradiation analyzed by a spot assay

The *S. elongatus* cultures of both treatments were adjusted to an optical density *OD*_750_ = 0.5, and a dilution series to the power of 10 was made in BG11 medium (10^0^ – 10 ^−5^). 5 µ*L* of each dilution was dropped on BG11-agar plates and cultivated at 28 °*C* ^33^ under constant light with the intensity of 30 µ*mol* photons s ^−1^ m ^−2^ for 7 days. All experiments are shown in Fig. 1D. Top: Non-irradiated control in a dilution series (1:10), three replicates. Fig. 1D below: Irradiated sample in a dilution series (1:10), three replicates. The bacteria were irradiated with a laser (*λ*_*ex*_ = 440 *nm*, power: 1.8 *mW*) for three minutes before preparation of a spot assay. Representative results are shown in Fig. 1D.

### The spectral properties of *S. elongatus*

To characterize the spectral properties of the photosynthetic pigments, absorption and emission spectra were recorded from 20 µ*L* of a cyanobacterial suspension, embedded in a low-melting agarose matrix to prevent cell movement (Fig. 1C). The absorption spectrum shown in Fig. 1B features four distinct bands: the soret band of chlorophyll a at 440 *nm* ^34^, the carotenoid band at 500 *nm* ^35^, the PBS band at 630 *nm* ^36^ and the *Q*_*y*_ band of chlorophyll a at 680 *nm* ^37^. Excitation of the soret band is very efficient, taking additional advantage of the large Stokes shift to separate the laser reflection at the cavity mirrors from the emission signal, which is dominated by the chlorophyll a emission at 680 *nm*. ^35^

## Acknowledgments

The authors would like to thank F. de Courcy for English proofreading of the manuscript, M. Kittelberger for initial experiments and M. Harter for the hint of trying quantum biology. Funding: A.J.M., F.W. and K.H. acknowledge support by the VW foundation (project title: A quantum beat for life) and the Deutsche Forschungsgemeinschaft (ME 1600/13-3; HA 2146/23-1; SFB 1101/D02, Z02).

## Author contributions

S.z.O.-K., K.H. and A.J.M. designed the project. T.R. and J.R. performed experiments. T.R. and F.W. analyzed the data and wrote the manuscript with input and proofreading from K.F., K.H., A.J.M., S.z.O.-K. and J.R.

## Data Availability

Data are available in the main text or the supplementary materials. Further material is available from the corresponding author upon reasonable request.

## Supplementary Materials

Materials and Methods

Supplementary Text

Figures S1-S2

References 1-7

