## Supplementary Materials for "Long-range quantum coherence of the photosystem 2 complexes in living cyanobacteria"

**This PDF file includes:**

Supplementary Text
Figs. S1 and S2
References

**Blank microcavity**

White light transmission spectra were used to determine the cavity resonance for different mirror separations as shown in Fig. S1. Here, the mirror distance of an empty microcavity was reduced by the piezo actuators from the top to the bottom and the spectral position of the cavity resonance was tuned from $\lambda = 710\,nm$ to $640\,nm$. This allows to tune the cavity resonance across the absorption and emission maximum of the cyanobacteria at $\lambda = 680\,nm$. The upper, movable mirror was slightly curved so that the distance between the two mirrors is minimal in the center and increases radially leading to so called Newton rings. Fluorescence spectra and fluorescence decay curves were recorded as a function of the mirror distance by focusing a $440\,nm$ laser beam with the same high NA objective lens into the cavity.

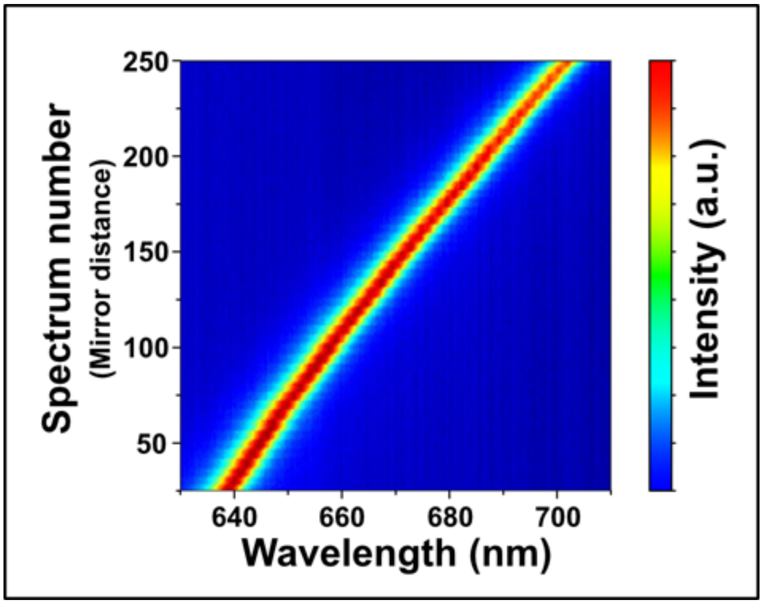


**Fig. S1:** White light transmission spectra of an empty microresonator as a function of the mirror distance. For each consecutive spectrum, the mirror spacing is increased, which shifts the resonance wavelength of the cavity accordingly.

**Time correlated single photon counting (TCSPC)**

Quantitative fluorescence intensity measurements of living cyanobacteria are not trivial, since the number of pigments of a particular bacterium varies or individual pigments of the photosystem can photo-bleach during an experiment. However, the fluorescence lifetime is independent of the concentration and can be determined by time correlated single photon counting (TCSPC).

The fluorescence lifetime $\tau$ is a measure of the average dwell time of molecules in the excited state. The fluorescence intensity decays exponentially, which can be described by [1]:

$$I(t) = I_0 e^{-t/\tau} \quad (1)$$

$t$: Time elapsed after excitation

$I(t)$: Fluorescence intensity after time t

$I_0$: Fluorescence intensity directly after excitation

Competitive processes such as internal conversion, thermal equilibration, intersystem crossing or energy transfers, which can take place instead of fluorescence, reduce the average fluorescence lifetime.

Usually, the fluorescence lifetime of organic fluorophores is in the lower nanosecond range. Since the bacteria need the harvested solar energy for biochemical reactions, the energy transfer to the reaction center must happen much faster. Due to the extremely fast energy transfer to the reaction center, the fluorescence lifetime decreases into the picosecond range. The fluorescence lifetime was acquired by TCSPC. The living bacteria are excited with very short laser pulses $(< 80\,ps)$ and a pulse rate of $80\,MHz$. The observed temporal decays are a convolution of the fluorescence decay with the instrument response function (IRF). The fluorescence lifetime can be determined through an iterative deconvolution of the IRF. Cyanobacteria have complicated decay properties due to their very complex structure. The decay must be represented by a sum of exponential functions [2]:

$$I(t) = \sum_i A_i\, e^{-t/\tau_i} \quad (2)$$

$A_i$ determines the amplitude or weighting of the individual exponents. From this equation the average intensity weighted fluorescence lifetimes $\tau_I$ can be determined [2]:

$$\tau_I = \frac{\sum_i A_i \, \tau_i^2}{\sum_i A_i \, \tau_i} \tag{3}$$

As shown in [2], $\tau_I$ is very accurate and it is largely independent of the sum of the exponential functions. It can, in free space, be regarded as an intrinsic property of the fluorophore. However, a cavity influences the spontaneous emission rate and thus the fluorescence lifetime. The probability of the spontaneous emission rate is higher if the emitter is inside a resonant cavity, i.e. Purcell effect [3].

**Mathematical description of two coupled harmonic oscillators**

In order to model strong coupling between the cavity and the bacteria we used two coupled damped harmonic oscillators. The differential equations of motion of a coupled oscillator system can be written as:

$$\ddot{x}_1(t) + \gamma_1 \dot{x}_1(t) + \omega_1^2 x_1(t) + \kappa x_2(t) = 0 \tag{4}$$

$$\ddot{x}_2(t) + \gamma_2 \dot{x}_2(t) + \omega_2^2 x_2(t) + \kappa x_1(t) = 0 \tag{5}$$

with the damping constants $\gamma_1, \gamma_2$, the resonance frequencies $\omega_1, \omega_2$ of the two oscillators and the coupling constant $\kappa$. The first oscillator $x_1$ is used to model the cavity mode, while the second oscillator $x_2$ describes the bacteria emission. Equation (4) and (5) are coupled via the terms proportional to $\kappa$. This allows an energy exchange between the cavity $x_1$ and the bacteria $x_2$. The temporal amplitudes $x_1(t)$ and $x_2(t)$ can be obtained by solving these equations numerically and the result for the special case of $\omega_1 = \omega_2 = 680\ nm$, $\gamma_1 = \gamma_2 = 0.03\ eV$ and $\kappa = 0.4\ eV$ is shown in Fig. S2.

The temporal response of the cavity $x_1(t)$ and of the bacteria $x_2(t)$ is presented in Fig. S2 A/B, respectively. The starting amplitude of the cavity is set to $x_1(0) = 1$ since it is directly excited. In this example, the bacteria is exclusively excited via strong coupling and $x_2(0)$ is zero. The energy exchange between the cavity and the bacteria leads to a beating pattern of the temporal amplitudes and the energy is transferred back and forth between the cavity and the bacteria. The temporal amplitudes $x_1(t)$, $x_1(t)$ can be seen as the autocorrelation functions of the coupled system and their Fourier transform is the corresponding spectrum, which is presented in Fig. S2C/D, respectively. Both the transmission spectrum of the cavity in C and the bacteria emission in D show two polaritonic modes, which are caused by strong coupling. This anti-crossing behavior for $\omega_1 = \omega_2$ is experimentally observed in Fig. 4 of the main text. There is an excellent agreement between the spectra calculated with this harmonic oscillator approach and the experimental data in Fig. 4.

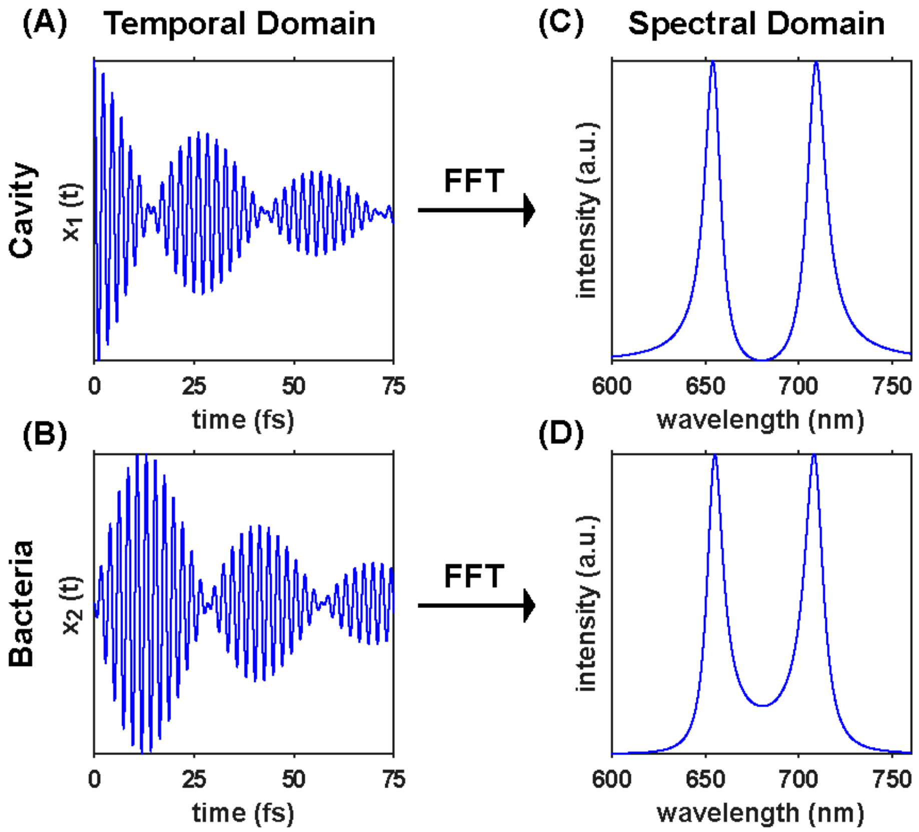


**Fig. S2: A/B** show the temporal amplitude of the two harmonic oscillators described equation in 4 and 2, where $x_1(t)$ describes the cavity mode and $x_2(t)$ is used to model the bacteria. The corresponding spectra are shown in **C/D** and are calculated by Fourier transformation of A/B.

**Calculation of the number of coupled chlorophyll a molecules participating in splitting.**

The mode volume ($V_m$) can be calculated according to Equation (6), where $d$ is the optical path length of the cavity and $r_0$ the radius of the excitation spot. [4]

$$V_m = \frac{\pi r_0^2 d}{4} \tag{6}$$

The transition dipole moment of chlorophyll a in BG11 medium with a refractive index of $n = 1.34$ is $5.39\,D$ [5]. Equation (7) was used to estimate the number of coupled chlorophyll a molecules in the excitation spot. [4]

$$N = \frac{(\Delta E^{vac}(N))^2}{\frac{2\mu_{12}^2 \hbar\omega}{n^2 \varepsilon_0 V_m}} \tag{7}$$

The experimental splitting before bleaching observed in Fig. 4D is 0.0478 eV, which is caused, according to equation (7), by approximately $4.8 \cdot 10^5$ coupled chlorophyll a molecules. After bleaching, the splitting was reduced to 0.0305 eV, which corresponds to approximately $2.0 \cdot 10^5$ fluorophores.

For comparison, the number of chlorophyll a molecules per bacterium is reported to $3 \cdot 10^4$ PS2 per bacterium [6] and $35$ chlorophyll a molecules per PS2 [7]. The excitation spot contains about $5.2 \cdot 10^5$ chlorophyll a molecules (only about half of a bacterium is irradiated). This number of chlorophyll a molecules should result in a theoretical splitting of 0.0333 eV. This number can only roughly be compared to the experimental splitting observed in this work, since the bacteria vary in size and contain different numbers of photosystems depending on the cultivation conditions. However, the expected theoretical splitting is in the same order of magnitude as our measurement.
